# Spatial Transcriptomics Reveals Compartment-Specific Immune Activation Signatures in Ileal and Lymph Node Tissue in Treated HIV Infection

**DOI:** 10.64898/2026.08.28.747869

**Authors:** Maura Barrett, Jodi Anderson, Kevin Escandón, Garritt Wieking, Ty Schroeder, Erik Swanson, Melanie L. Graham, Michael Gale, Timothy W. Schacker, Nichole R. Klatt, Christopher M. Basting

## Abstract

People with HIV (PWH) on long-term antiretroviral therapy (ART) continue to experience elevated rates of morbidities and mortality driven by persistent immune activation despite viral suppression. Known contributors include low-level HIV provirus activity, microbial translocation in part from epithelial barrier dysfunction, microbiome dysfunction, and co-infections. However, how these interact and where they predominate across tissue compartments remains incompletely defined. Here, we applied spatial transcriptomics to characterize compartment-specific transcriptional programs in ileum (epithelium, Peyer’s patches, lamina propria) and inguinal lymph nodes (B Cell follicles and T cell zone) from ten PWH on long-term ART, stratified by CD4/CD8 ratio into low-ratio and high-ratio groups, with low-ratio as a proxy for immune activation and increased risk for non-AIDS related serious event. Comparison of global expression found significant differences between groups in four of five compartments. Differential expression analysis identified 483 differentially expressed genes across four of five compartments, with the greatest burden in the T-cell zone and none in the lamina propria. Gene set enrichment analysis identified 116 enriched pathways predominantly in the low-ratio group, spanning immune activation, infection-associated, and metabolic programs, with Peyer’s patches showing the broadest transcriptional divergence of any compartment. Cross-compartment signals included higher expression of ORMDL3 and ARL17B in the low-ratio group implicating mitochondrial stress and inflammasome activation, lower expression of CCL3L3 and FCMR in the low-ratio group suggesting impaired immune execution, and divergent ribosomal protein programs between B-cell follicles and the T-cell zone. Cell deconvolution identified compartment-specific differences in estimated immune cell proportions, and T-cell zone gene expression showed significant associations with HIV reservoir measures and plasma markers of microbial translocation and immune activation. Together these findings support spatially heterogeneous immune activation as a feature of persistent immune dysregulation in treated HIV infection and provide compartment-resolved, hypothesis-generating evidence for the tissue-specific mechanisms driving inflammation in this population.

## Introduction

Antiretroviral therapy (ART) has markedly improved outcomes for people with HIV (PWH), reducing plasma viral loads to undetectable levels and enabling near-normal life expectancy (1–5). Yet, PWH on ART continue to experience higher rates of morbidities and mortality relative to the general population, a disparity driven in large part by chronic inflammation and immune activation despite viral suppression (3,6–9). Elevated markers of immune activation and inflammation are consistently associated with adverse clinical outcomes, including cardiovascular disease, neurocognitive decline, and all-cause mortality, and these risks are substantially higher in PWH with inverted CD4/CD8 ratios, reflecting increased immune dysregulation that no current therapy adequately resolves (3,9–17).

Persistent immune activation in PWH on ART remains incompletely understood and is likely multifactorial (3,8,9,13). Proposed contributors include low-level transcription and translation of HIV provirus gene products, epithelial barrier dysfunction and microbiome dysbiosis that results in microbial translocation, and co-infections (3,8,9,13,14). Low-level viral activity may sustain immune responses through ongoing antigenic stimulation, while epithelial barrier damage increases gut permeability, allowing microbial products to enter circulation and trigger innate immune activation (3,8,13,14). Co-infections, including reactivation of latent pathogens such as cytomegalovirus (CMV) and Epstein-Barr virus (EBV), provide additional inflammatory stimuli that further amplify immune activation (3,8,13,14). These mechanisms likely act in combination, with recent evidence suggesting bidirectional relationships between HIV provirus gene products, systemic cytokines, T cell activation, and loss of commensal microbes in the gastrointestinal tract (3,13,18–22). However, the relative contributions of these processes to chronic immune activation, and the tissue compartments in which they predominate, remain incompletely defined.

Systemic biomarkers such as interleukin (IL)-6 and C-reactive protein (CRP) have been associated with immune activation and clinical outcomes despite ART, but provide limited insight into the tissue-specific and microenvironmental sources driving this inflammation (23). Similarly, traditional transcriptomic approaches, including bulk and single-cell sequencing, have provided important insights into HIV-associated immune dysfunction but cannot resolve spatial organization or microenvironmental context (24,25). Bulk analyses average signals across heterogeneous cell populations, while single-cell approaches disrupt tissue architecture, obscuring compartment-specific signals (24,25). In contrast, spatially resolved approaches preserve tissue structure and enable identification of localized gene expression patterns (24–26). This is particularly critical in HIV infection, where immune activation, viral persistence, and immune regulation occur within non-uniform anatomical niches, and where integrating transcriptomic and microenvironmental data can reveal features not detectable with non-spatial approaches (12,16,24,25,27,28).

Spatial resolution is especially important in the gastrointestinal (GI) tract and lymphoid tissues (15,17,27–30). The GI tract contains the majority of the body’s immune cells and serves as a major HIV reservoir and interface between host immunity and the microbiome (14,17). Gut-associated lymphoid tissue (GALT) comprises distinct compartments, including the epithelium, lamina propria, and Peyer’s patches, that are differentially affected during infection (15,16,31–36). Disruption of these compartments contributes to microbial translocation and sustained immune activation, while uneven antiretroviral drug distribution may permit localized viral activity (13,14,17,27). Lymph nodes are equally critical sites of HIV pathogenesis, where viral reservoirs accumulate within follicular dendritic cell networks and B cell follicles that function as microanatomical “sanctuaries” for infected cells (29,30). Antiretroviral drug penetration is also variable across lymphoid compartments, with reduced concentrations in lymph nodes compared to blood, potentially contributing to suboptimal tissue-level suppression of virus (27–30). Given the severe clinical consequences and the incomplete understanding of mechanisms driving persistent immune activation in PWH, elucidating signatures of it in the GI tract and lymph nodes represents a critical area of investigation (29).

Here, we utilized spatial transcriptomics to characterize compartment-specific transcriptional programs within ileum and inguinal lymph node tissue, including Peyer’s patches, lamina propria, and epithelium in ileal tissue and B cell follicles and T cell zones in the lymph nodes. Tissue samples were from PWH on long-term ART (>2 years) with undetectable plasma viral loads and were stratified by their CD4+/CD8+ ratios into high-ratio and low-ratio groups, a distinction previously shown to be associated with immune activation and mortality risk in PWH(37,38). Through this approach, we sought to define molecular and transcriptional signatures associated with persistent immune activation at the compartment level to gain an understanding of the tissue specific mechanisms driving inflammation in treated HIV infection.

## Results

Table 1 enumerates average baseline characteristics of the two groups, all participants were male. Observed differences between groups were consistent with their CD4/CD8 ratio stratification. Low-ratio individuals were older on average (53 vs. 31 years) and had longer HIV infection duration (15 vs. 4 years). The low-ratio group had lower absolute CD4+ T cell counts (511 vs. 716 cells/μL), higher absolute CD8+ T cell counts (1,410 vs. 556 cells/μL), and a lower CD4/CD8 ratio (0.36 vs. 1.29) compared to the high-ratio group. BMI was similar between groups (26.3 vs. 25.8 kg/m²).

**Table 1.** Baseline characteristics of enrolled participants.

|  | Low CD4/CD8<br>Ratio (n = 5) | High CD4/CD8<br>Ratio (n = 5) | p-value* |
| --- | --- | --- | --- |
| **Characteristic** |  |  |  |
| Age (years) | 53 (7.7) | 29 (7.9) | 0.001 |
| BMI (kg/m<sup>2</sup>) | 26.3 (1.9) | 25.4 (3.7) | 0.641 |
| Time living with HIV (years) | 15 (9.5) | 4 (5.5) | 0.062 |
| Absolute CD4<sup>+</sup> T cell count<br>(cells/mm<sup>3</sup>) | 511 (215.5) | 716 (237.2) | 0.192 |
| Absolute CD8<sup>+</sup> T cell count<br>(cells/mm<sup>3</sup>) | 1410 (477.4) | 556 (178.2) | 0.013 |
| CD4/CD8 ratio | 0.36 (0.1) | 1.29 (0.2) | < 0.001 |

Clinical characteristics of participants, all of whom are male. Variables presented as mean (standard deviation).
\* Welch Two Sample t-test
Participants (n = 10; all male) were stratified into Low CD4 Ratio (n = 5) and High CD4 Ratio (n = 5) groups. Values are presented as mean (SD). Age, time living with HIV, absolute CD4<sup>+</sup> count, and absolute CD8<sup>+</sup> count are rounded to the nearest whole number. Body mass index (BMI) and all standard deviations are rounded to one decimal place, and the CD4 ratio is reported to two decimal places. Between-group comparisons were performed using unpaired two-tailed Welch's t-tests. P-values are reported to three decimal places, with values < 0.001 reported as "<0.001." A p-value < 0.05 was considered statistically significant.

### Principal component analysis shows distinct separation of CD4/CD8 ratio groups across tissue compartments

To assess whether global gene expression profiles differed between low- and high-ratio groups within each compartment, principal component analysis (PCA) was performed on expression data. Varying degrees of group separation was seen in the first two principal components (Fig. 1). To formally test multivariate differences, patient-level PERMANOVA was performed on each expression profile. In the ileum, Peyer’s patches showed the strongest group separation, with PC1 explaining 78.9% of variance and significant multivariate separation by PERMANOVA (R² = 0.942, p = 0.03). Epithelium similarly showed distinct separation (PC1: 54.0%, R² = 0.70, p = 0.03). The lamina propria was the exception, with modest PC1 variance (18.9%) and non-significant group separation (R² = 0.226, p = 0.059). In the lymph nodes, both B-cell follicles (PC1: 58.3%, R² = 0.712, p = 0.027) and T-cell zones (PC1: 29.6%, R² = 0.558, p = 0.032) showed significant group separation. Overall, these findings demonstrate that global gene expression profiles are broadly remodeled across gut and lymph node tissue compartments in PWH with low versus high CD4/CD8 ratios, with the ileal lamina propria as the sole exception.

**Figure 1:**
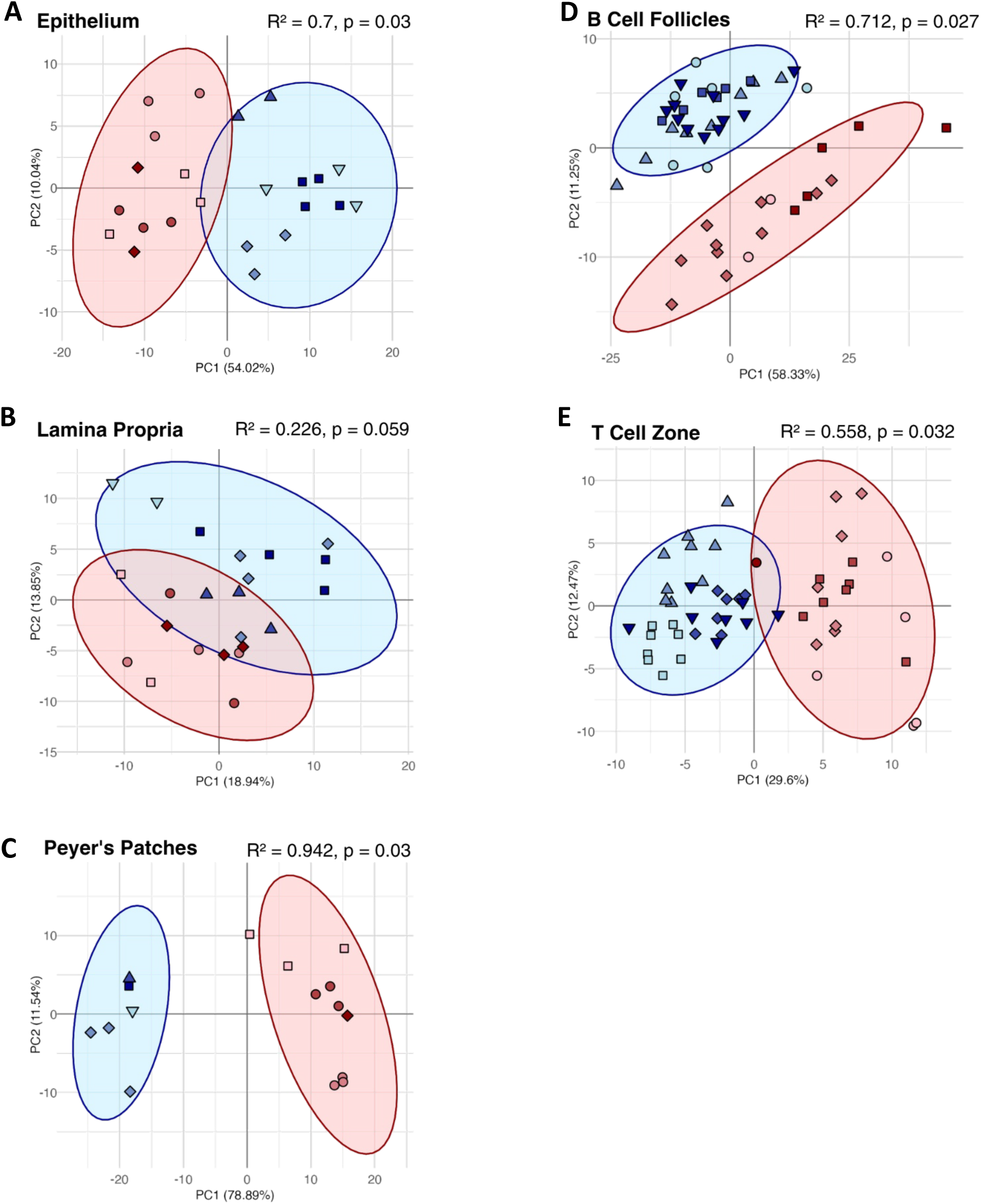
Principal Component Analysis of CD4/CD8 ratio groups across tissue compartments. PCA was performed on RUV4-normalized expression data for each compartment independently. Plots A–E show PC1 versus PC2 for the epithelium (A), lamina propria (B), Peyer’s patches (C), B-cell follicles (D), and T-cell zone (E). Each point represents one region of interest with shape differentiating patient ID (PID). Low-ratio individuals denoted in blue and high-ratio individuals in red. Shaded ellipses represent 95% confidence intervals for each group. PERMANOVA R² and p values and the percentage of variance explained by each principal component are shown on each plot.

**Figure 2.**
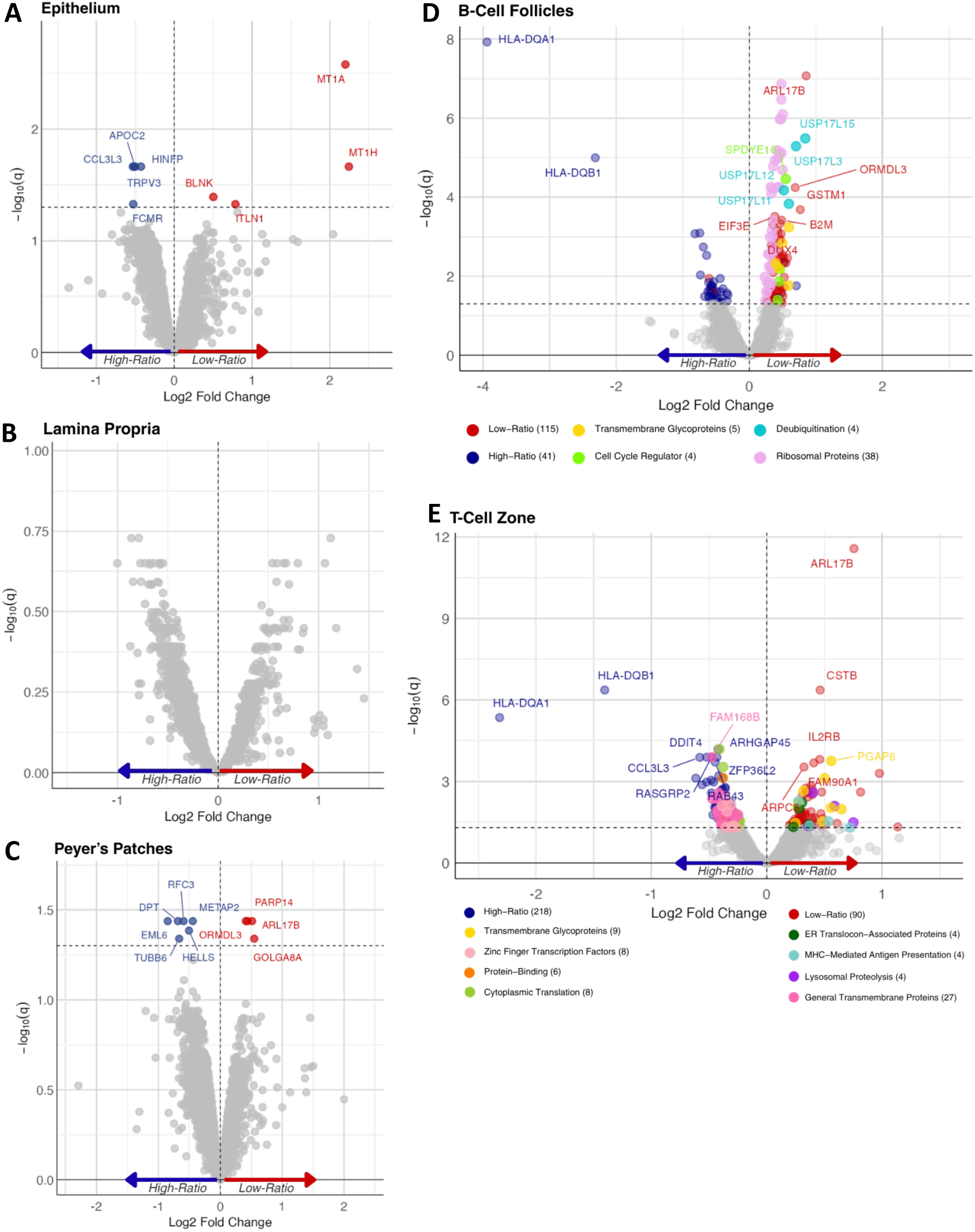
Differential gene expression between low- and high-ratio groups across tissue compartments. Volcano plots display differentially expressed genes (DEGs) for the epithelium (A), lamina propria (B), Peyer’s patches (C), B-cell follicles (D), and T-cell zone (E). The x- axis shows Log2 fold change (log2FC), where positive values indicate higher expression in the low-ratio group and negative values indicate higher expression in the high-ratio group. The y-axis shows -log10(q), where q is the FDR-adjusted p-value. The dashed horizontal line indicates the significance threshold (q = 0.05) and the dashed vertical line indicates log2FC = 0. In panels A–C, significantly upregulated genes (q ≤ 0.05) are shown in red and downregulated genes in blue, with non-significant genes in gray. D-E Gene labels are shown for genes with q ≤ 0.0005, excluding ribosomal protein subunit genes. Points are colored by functional cluster as indicated in the legend below each panel, with background genes and unclustered genes shown at reduced opacity. Low-ratio enriched and high-ratio enriched cluster labels reflect the directionality of expression within each cluster.

### Differential expression analysis reveals compartment-specific gene expression differences between CD4/CD8 ratio groups

Next, we identified differentially expressed genes (DEGs) between the low- and high-ratio groups, identifying 483 DEGs with FDR adjusted q-values ≤ 0.05 (9 in epithelium, 10 in Peyer’s patches, 0 in lamina propria, 156 in B-cell follicles, 308 in the T-cell zone), corresponding to 452 unique genes. Of these, 193 DEGs were upregulated in the low-ratio group (positive log_2_FC), while 259 were upregulated in the high-ratio group (negative log_2_FC). Due to the volume of DEGs in the lymph node compartments, the DAVID (Database for Annotation, Visualization, and Integrated Discovery) Gene Functional Classification Tool was used to group genes into functionally related clusters for each ratio group. Cluster labels were assigned descriptively based on predominant gene functions and enriched annotation terms reported by DAVID. Gene functions and protein annotations referenced throughout were obtained from NCBI Gene and GeneCards (39–41).

Nine genes with significant expression differences were identified in the epithelium (Fig. 1A), including 4 upregulated in the low-ratio group and five in the high-ratio group. The most significantly upregulated genes in the low ratio group were the metallothioneins MT1A (log_2_FC = 2.20, q = 0.003) and MT1H (log_2_FC = 2.25, q = 0.022), both involved in heavy metal binding and cellular stress responses, alongside ITLN1 (log_2_FC = 0.79, q = 0.047), a lectin involved in innate immunity, and BLNK (log_2_FC = 0.50, q = 0.040), a B-cell receptor signaling adaptor. Downregulated genes included CCL3L3 (log_2_FC = -0.53, q = 0.022), a CCR5-binding chemokine, FCMR/Toso (log_2_FC = -0.53, q = 0.047), an Fc receptor for IgM, TRPV3 (log_2_FC = -0.51, q = 0.022), a thermosensitive ion channel, HINFP (log_2_FC = -0.43, q = 0.022), a transcription factor involved in cell cycle regulation, and APOC2 (log_2_FC = -0.50, q = 0.022), an apolipoprotein involved in lipid metabolism.

In Peyer’s patches, 10 DEGs were identified (Fig. 1C). Upregulated genes in the low-ratio group included GOLGA8A (log_2_FC = 0.55, q = 0.046) and ARL17B (log_2_FC = 0.51, q = 0.037), involved in Golgi structure and vesicle-mediated transport, PARP14 (log_2_FC = 0.44, q = 0.037), a poly(ADP-ribose) polymerase with anti-apoptotic functions, and ORMDL3 (log_2_FC = 0.41, q = 0.037), an endoplasmic reticulum protein. Six genes were upregulated in the high-ratio group: EML6 (log_2_FC = -0.85, q = 0.037) and TUBB6 (log_2_FC = -0.66, q = 0.046), encoding microtubule-associated proteins, DPT (log_2_FC = - 0.68, q = 0.037), RFC3 (log_2_FC = -0.59, q = 0.037), required for DNA replication and cell cycle progression, METAP2 (log_2_FC = -0.44, q = 0.037), a regulator of protein synthesis, and HELLS (log_2_FC = -0.50, q = 0.041), a lymphoid-specific helicase implicated in cellular proliferation and lymphocyte development.

Differential expression analysis identified 156 DEGs in B-cell follicles, with 115 upregulated in the low-ratio group and 41 in the high-ratio group (Fig. 1D). DAVID gene functional classification identified four clusters among low-ratio upregulated genes totaling 51 genes: (1) Ribosomal Proteins, 38 genes; (2) Deubiquitination, 4 genes; (3) Cell Cycle Regulators, 4 genes; and (4) Transmembrane Glycoproteins, 5 genes. No clusters were identified in the high-ratio group.

The ribosomal protein cluster was dominated by 22 RPL and 15 RPS subunit genes, with PCBP1 (log_2_FC = 0.35, q = 0.006), an RNA-binding protein involved in mRNA stability and translation, as the only non-ribosomal member. The deubiquitination and cell cycle clusters each comprised four genes of the same family - USP17 genes associated with protein stability and apoptotic regulation, and SPDYE genes linked to cell-cycle regulation and protein kinase binding, respectively. The transmembrane glycoprotein cluster included CD164 (log_2_FC = 0.40, q = 0.005), a regulator of proliferation, adhesion, and migration of hematopoietic progenitor cells (HPCs), and EVI2B (log_2_FC = 0.50, q = 0.002), primarily expressed in HPCs and crucial for granulocyte differentiation.

In the T-cell zone, 308 DEGs were identified between low- and high-ratio groups (218 vs 90) (Fig. 1D). Functional gene classification identified four clusters in both the low- and high-ratio enriched gene lists. In the low-ratio group, these clusters were annotated as (1) Lysosomal Proteolysis, 4 genes; (2) MHC-Mediated Antigen Presentation, 4 genes; (3) ER Translocon-Associated Proteins, 4 genes; and (4) Transmembrane Glycoproteins, 9 genes.

The lysosomal proteolysis cluster comprised three cathepsin-family proteases and LGMN. CTSB (log_2_FC = 0.589, q = 0.008), CTSS (log_2_FC = 0.40, q = 0.003), and CTSD (log_2_FC = 0.36, q = 0.044) are involved in lysosomal degradation, antigen processing, autophagy, and apoptosis. LGMN (log_2_FC = 0.75, q = 0.032), a cysteine protease activated under acidic lysosomal conditions, contributes to processing bacterial peptides and endogenous proteins for MHC class II presentation. The MHC-mediated antigen presentation cluster spanned both class I and class II molecules: B2M (log_2_FC = 0.49, q < 0.001), essential for MHC class I surface stability, alongside HLA-DPA1 (log_2_FC = 0.53, q = 0.029) and HLA-DQA2 (log_2_FC = 0.72, q = 0.050), involved in antigenic peptide presentation, and HLA-DMA (log_2_FC = 0.37, q = 0.042), which supports peptide loading onto class II molecules. The ER translocon-associated cluster comprised genes involved in ER membrane translocation, glycosylation of nascent polypeptides, signal peptide cleavage, and proteasomal targeting; notably SPCS1 (log_2_FC = 0.31, q = 0.006), predicted to be involved in signal peptide processing, viral protein processing, and virion assembly.

The high-ratio group also yielded four clusters with generally more genes per cluster: (1) Protein-Binding, 6 genes; (2) Cytoplasmic Translation, 8 genes; (3) Zinc Finger Transcription Factors, 8 genes; and (4) General Transmembrane Proteins, 27 genes. The protein-binding cluster comprised six genes united by shared protein-binding activity but otherwise representing heterogeneous intracellular functions including mitochondrial homeostasis and mRNA post-transcriptional regulation, alongside several functionally uncharacterized members. The cytoplasmic translation cluster comprised four RPL and four RPS genes, distinct from those identified in the low-ratio group of B-cell follicles. Within the zinc finger transcription factor cluster, IKZF1 (log_2_FC = -0.39, q = 0.008) and KLF13 (log_2_FC = -0.31, q = 0.007) regulate lymphocyte differentiation and function respectively. The general transmembrane protein cluster, with 27 genes, was the largest; CD3G (log_2_FC = -0.29, q = 0.018) couples antigen recognition to T-cell signaling, LIME1 (log_2_FC = -0.35, q = 0.008) links receptor stimulation to downstream pathways, and TMIGD2 (log_2_FC = -0.42, q = 0.019) positively regulates T-cell activation and cytokine production. CD6 (log_2_FC = -0.32, q = 0.013) and CD96 (log_2_FC = -0.34, q = 0.013) support adhesive interactions of activated T and NK cells, with CD6 additionally promoting continuation of T-cell activation. SIRPG (log_2_FC = -0.26, q = 0.021) and SIGIRR (log_2_FC = -0.29, q = 0.013) negatively regulate distinct immune signaling pathways.

### Gene set enrichment analysis identifies shared and compartment-specific pathway enrichment predominantly in low-ratio individuals

We next performed gene set enrichment analysis (GSEA) to identify pathway-level differences between low- and high-ratio groups across compartments (FDR q ≤ 0.05). In total 116 enriched pathways (76 unique) were identified from four compartments (57 in Peyer’s patches, 41 in B-cell follicles, 13 in the lamina propria, and 5 in T-cell zones). Only one pathway was enriched in the high-ratio group: kinesin motor pathways in Peyer’s patches, and 0 pathways were found in the epithelium. Pathways stemmed from 11 Reactome parent categories. Disease and Immune System (n = 20 each), followed by Metabolism of Proteins (n = 15), were the three largest categories. Figures 3 and 4 summarize normalized enrichment scores (NES), adjusted significance, parent categories, and cross-compartment patterns of enriched pathways, highlighting both shared pathways and compartment-specific programs.

**Figure 3.**
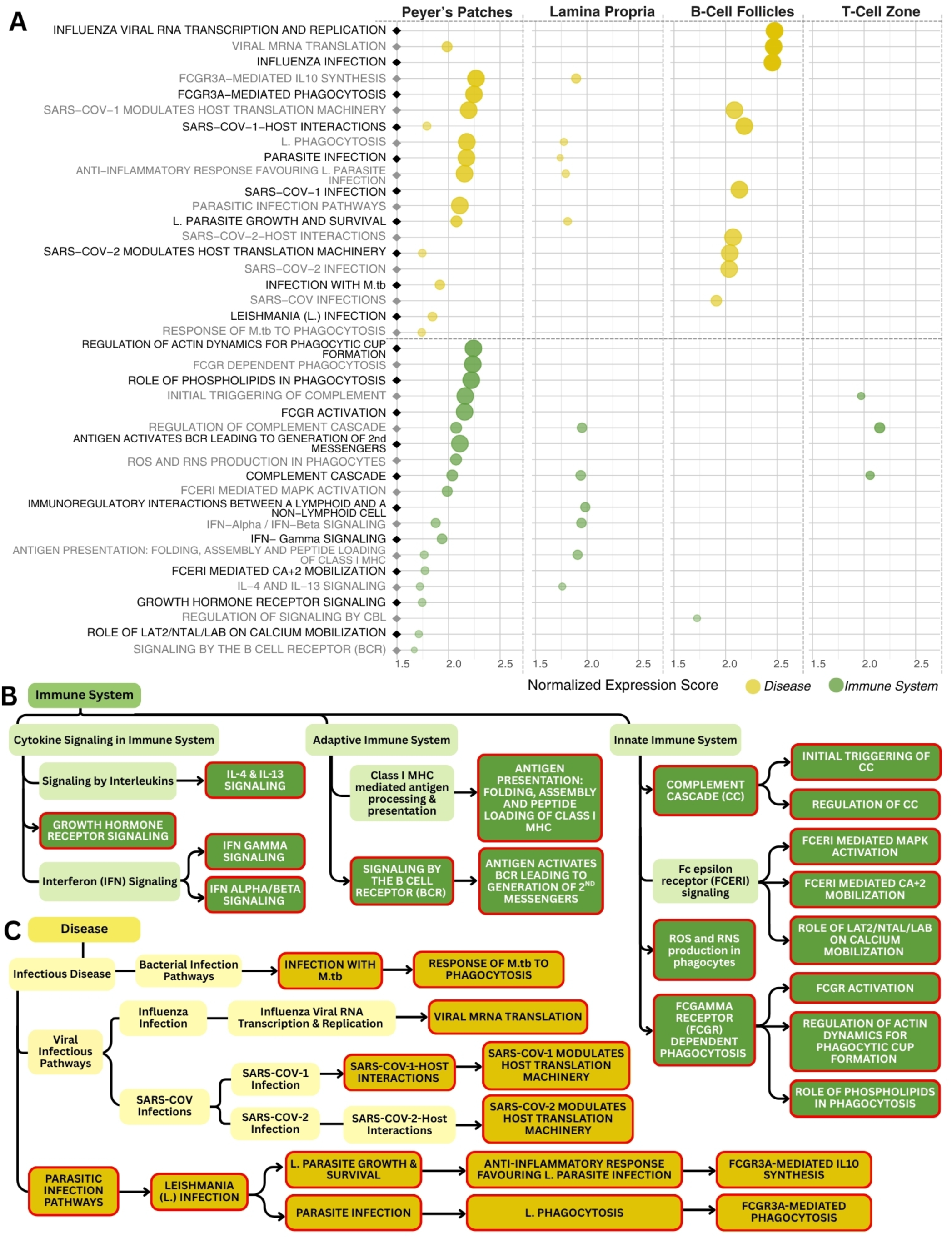
Disease and immune pathway enrichment across tissue compartments. GSEA identified enriched Reactome pathways in the low-ratio group across four compartments (FDR q ≤ 0.05); kinesin motor pathways in Peyer’s patches was the sole high-ratio enriched pathway. Panel A shows an S-plot of disease and immune system pathways across all four compartments, with each point representing one pathway in one compartment, colored by Reactome parent category, point size reflecting - log_10_(adjusted p-value), and opacity reflecting NES. Panel B shows the Reactome hierarchical tree for immune system pathways in Peyer’s patches; darker shading indicates significance and red outlines indicate low-ratio enrichment. Panel C shows the equivalent hierarchical tree for disease pathways in Peyer’s patches using the same conventions as Panel B.

**Figure 4.**
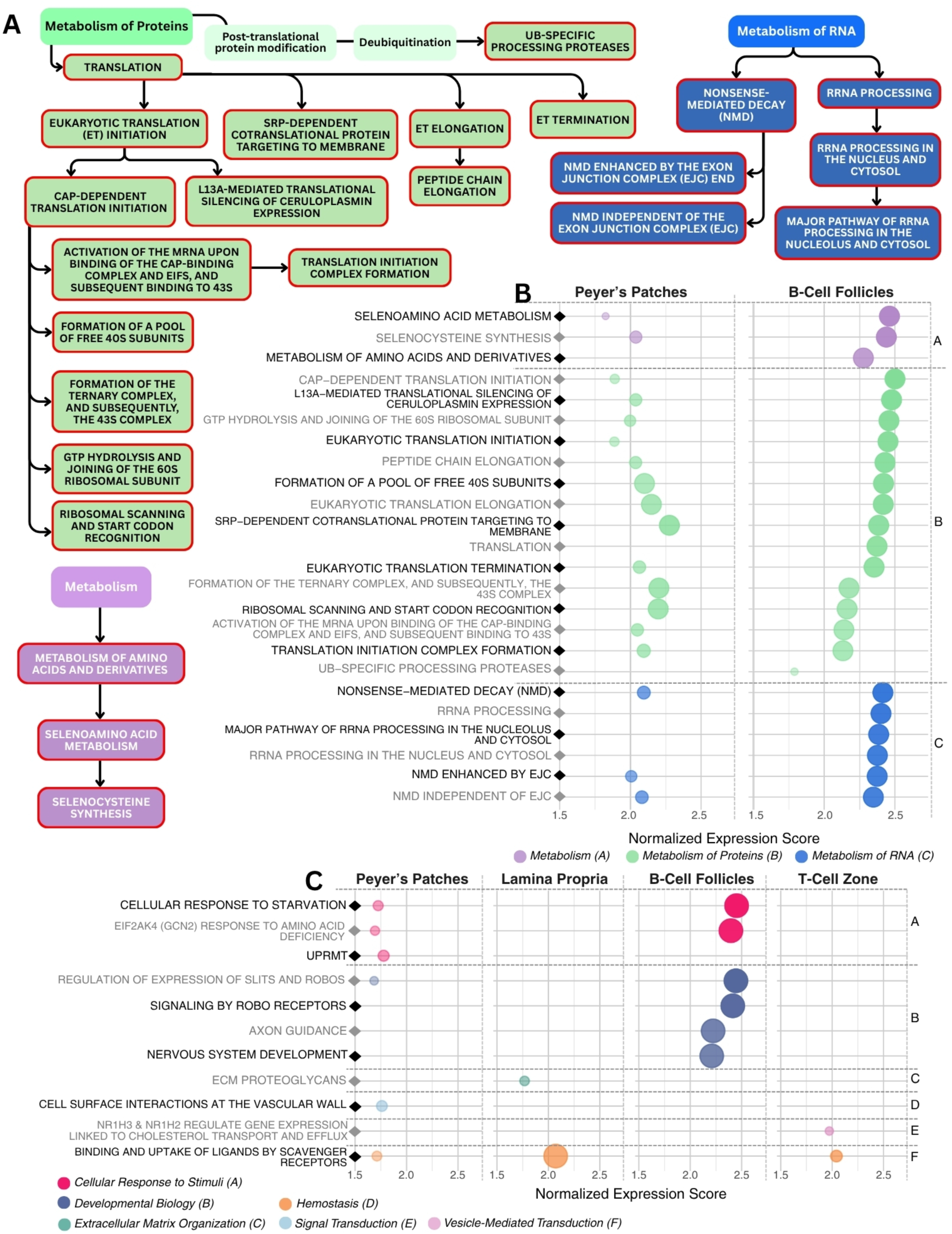
Metabolic and additional pathway enrichment across tissue compartments. GSEA identified enriched Reactome pathways in the low-ratio group across four compartments (FDR q ≤ 0.05). Panel A shows the Reactome hierarchical tree for metabolism categories in B-cell follicles, spanning Metabolism of Proteins, Metabolism of RNA, and Metabolism parent categories; darker shading indicates significance and red outlines indicate low-ratio enrichment. Panels B and C show S-plots of metabolism pathways and remaining parent categories respectively across all compartments, with each point representing one pathway in one compartment, colored by Reactome parent category, point size reflecting -log_10_(adjusted p-value), and opacity reflecting NES. Parent category colors are consistent across panels.

Disease-related pathways in the low-ratio group were found in Peyer’s patches (n = 14), B-cell follicles (n = 10), and the lamina propria (n = 5), which were further classified as parasitic, bacterial, or viral infection processes. Leishmania-associated pathways, including anti-inflammatory and phagocytic responses, were present in Peyer’s patches and the lamina propria, whereas mycobacterium tuberculosis (M.tb) related pathways were restricted to Peyer’s patches. Viral pathways were observed in Peyer’s patches and B-cell follicles, including influenza and SARS-CoV-1/2 modules that converged on viral mRNA translation and modulation of host translational machinery.

Immune system pathways were enriched most broadly in Peyer’s patches (n = 18), with fewer in the lamina propria (n = 6), T-cell zones (n = 3), and B-cell follicles (n = 1). Complement cascade initiation and regulation, seen in Peyer’s patches, the lamina propria, and T-cell zones, were the most broadly shared innate pathways. Additional innate (Fc receptor signaling, phagocytosis, ROS/RNS), cytokine (IFN-α/β, IFN-γ, IL- 4/IL-13), and adaptive (BCR signaling, MHC class I antigen presentation) pathways were enriched in Peyer’s patches. All immune pathways enriched in the lamina propria were also enriched in Peyer’s patches, with the exception of Immunoregulatory interactions between a Lymphoid and a non-Lymphoid cell (NES = 1.98, q = 0.007), which was unique to the compartment; regulation of signaling by CBL (NES = 1.72, q = 0.040) was the only immune pathway enriched in B-cell follicles.

Metabolic enrichment, from parent categories Metabolism of Proteins, Metabolism of RNA, or Metabolism, was localized to Peyer’s patches and B-cell follicles (Fig. 4). Protein-related enrichment was characterized by eukaryotic translation pathways (initiation, elongation, termination). RNA-related pathways centered on nonsense-mediated decay (NMD); B-cell follicles also had enrichment across the rRNA processing within the Metabolism category (Peyer’s patches: NES = 2.04, q < 0.001; B-cell follicles: NES = 2.44, q < 0.0001).

Figure 4 highlights enrichment within the remaining parent categories. EIF2AK4 (GCN2)-mediated nutrient sensing (Cellular Responses to Stimuli) was enriched in Peyer’s patches (NES = 2.40, q < 0.0001) and B-cell follicles (NES = 1.69, q = 0.040). Scavenger receptor-mediated ligand uptake, under Vesicle-Mediated Transport, appeared across Peyer’s patches (NES = 1.71, q = 0.036), the lamina propria (NES = 2.07, q < 0.0001), and T-cell zones (NES = 2.04, q = 0.017). This pathway co-occurred with a vascular wall interaction pathway in Peyer’s patches and ECM proteoglycan enrichment in the lamina propria (NES = 1.77, q = 0.037). Other compartment-specific paths included one or more pathways within Cellular Response to Stimuli in Peyer’s patches, Developmental Biology in B-cell follicles, and Signal Transduction in T-cell zones.

### T-cell zone gene expression correlates with HIV reservoir measures and systemic immune activation markers

Spearman correlations were performed between DEGs in each compartment and matching HIV reservoir measures in lymph nodes (viral DNA, viral RNA), as well as plasma biomarkers (IL6, sCD14, LBP, and I-FABP). Significant associations (nominal p- value ≤ 0.05, exploratory FDR-adjusted q ≤ 0.25) were identified exclusively for genes within the T-cell zone compartment (Fig. 5). Eight genes were positively correlated with HIV DNA and 23 genes were negatively correlated with HIV RNA; all were enriched in the high-ratio group. One gene inversely associated with HIV RNA, SERINC5 (ρ = - 0.88, p = 0.004, q = 0.192), is a host restriction factor shown to impair viral membrane fusion and entry (42–44). The third plasma biomarker with significant correlations was sCD14, which had positive (12) and negative (11) associations. Positive correlations were found exclusively with genes enriched in the low-ratio group, such as STMN1 (ρ = 0.86, p = 0.007, q = 0.224), a regulator of microtubule dynamics linked to cellular activation and cytoskeletal remodeling (45). In contrast, negative associations were observed with genes enriched in the high-ratio group, including FCGBP (ρ = -0.90, p = 0.002, q = 0.161), a glycoprotein linked to barrier-associated immune defense and pathogen exclusion (46).

**Figure 5.**
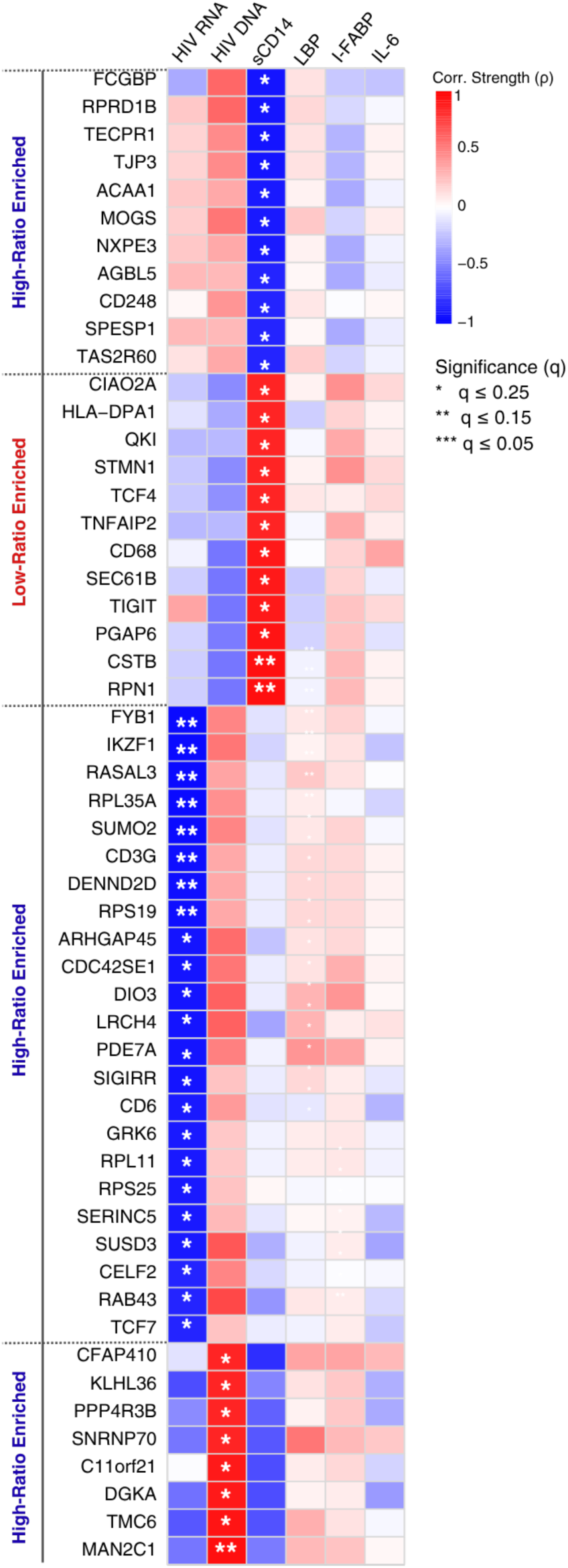
Spearman correlations between T-cell zone differentially expressed genes and biomarkers. Heatmap showing Spearman correlation coefficients between T-cell zone DEGs with at least one significant association and six external biomarkers: HIV RNA and DNA measured by in situ hybridization in lymph node tissue, and plasma markers of microbial translocation and immune activation (sCD14, LBP, I-FABP, and IL-6). Only DEGs with at least one significant association (nominal p ≤ 0.05, exploratory FDR q ≤ 0.25) are shown. Color indicates the direction and magnitude of correlation, ranging from blue (negative) to red (positive). Asterisks indicate significance thresholds: * q ≤ 0.25, ** q ≤ 0.15, *** q ≤ 0.05.

### Cell deconvolution identifies compartment-specific differences in estimated immune cell proportions between low- and high-ratio individuals

Cell deconvolution was performed using SpatialDecon to estimate proportions of immune cell types within each compartment (Fig. 6A) and assess differences between low- and high-ratio groups at a significance threshold of p ≤ 0.05 and exploratory FDR q ≤ 0.25 (Fig. 6B-D). In the epithelial compartment, mean estimated endothelial cell fractions were significantly higher in the low-ratio group compared to the high-ratio group (p < 0.0001, q < 0.001), with proportions of 0.03 vs. 0.02, corresponding to a 23.39% percent difference (PD). In the Peyer’s patches, three immune populations were increased in low-ratio individuals: neutrophils (p = 0.021, q = 0.137, 17.22% PD), natural killer (NK) cells (p = 0.012, q = 0.137, 22.08% PD), and CD4 memory T cells (p = 0.023, q = 0.137, 43.45% PD). These percent differences corresponded to modest shifts in estimated proportions between groups (Fig. 5C). CD4 naive T cells were increased in high-ratio individuals (p = 0.035, q = 0.158, 53.85% PD), mean estimated proportions were 0.14 (low-ratio) and 0.25 (high-ratio).

**Figure 6.**
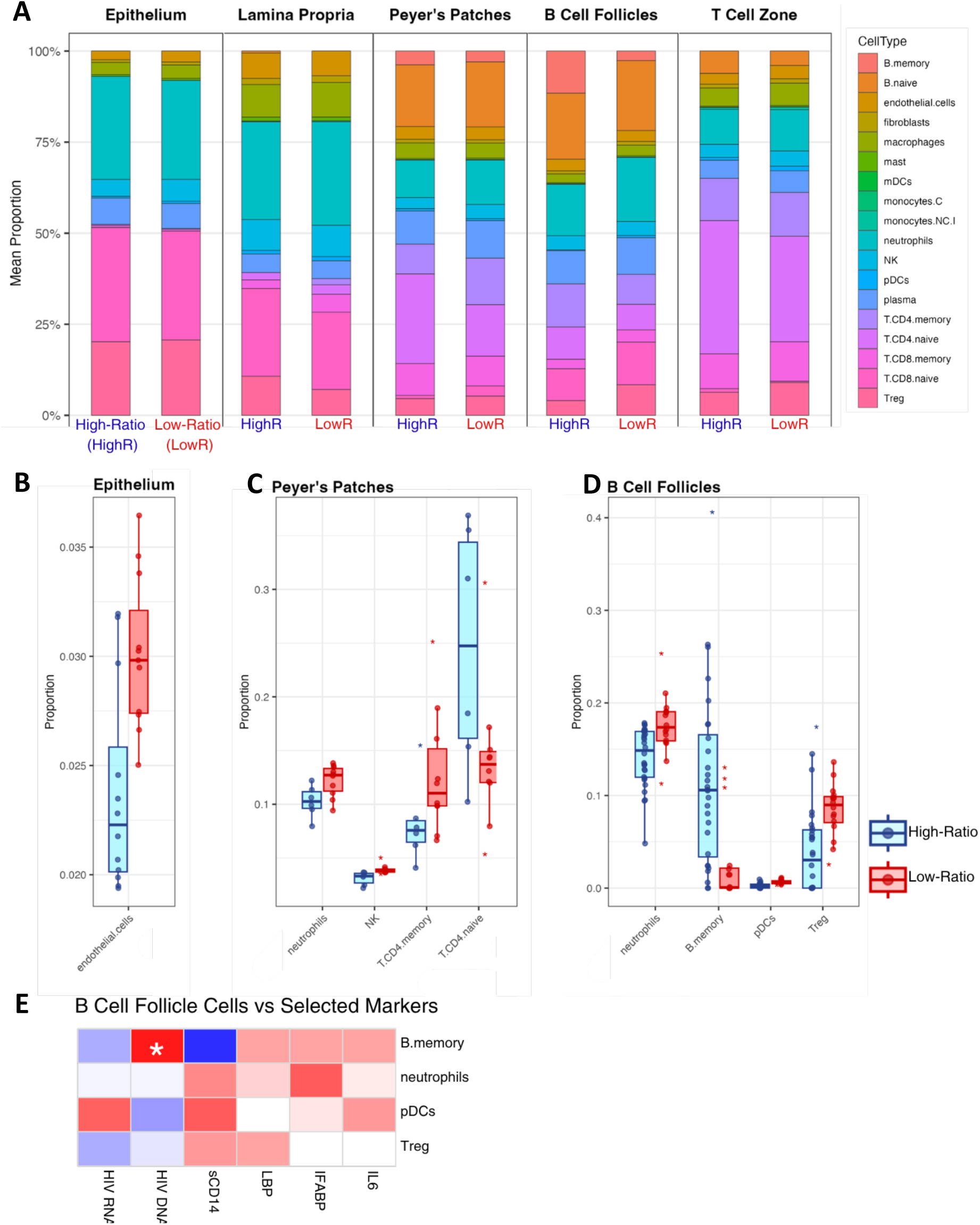
Estimated immune cell type composition and significant differences between low- and high-ratio groups. Cell type proportions were estimated using SpatialDecon with the safeTME reference matrix. Panel A shows mean estimated cell type composition as stacked bar plots across all five compartments (epithelium, Peyer’s patches, lamina propria, B-cell follicles, and T-cell zone), with each bar representing the mean proportions across all individuals in each group. Panels B-D show box plots of cell type proportions for cell populations with statistically significant differences between groups (nominal p ≤ 0.05, exploratory FDR q ≤ 0.25) in the epithelium (B), Peyer’s patches (C), and B-cell follicles (D). Low-ratio individuals are shown in red and high-ratio individuals in blue. Individual points represent per-patient estimated proportions; asterisks indicate outlier values. Panel E shows the Spearman correlation between B-memory cell abundance and HIV DNA in B-cell follicles, the only significant association identified between estimated cell type proportions and immune activation markers.

In B-cell follicles, 3 cell populations were increased in low-ratio individuals; neutrophils (p = 0.013, q = 0.122, 21.80% PD), plasmacytoid dendritic cells (pDCs; p = 0.014, q = 0.122, 79.78% PD), and regulatory T cells (Tregs; p = 0.041, q = 0.182, 70.07% PD). B-memory cells were increased in high-ratio individuals (p = 0.020, q = 0.122, 125.73% PD), with estimated proportions of 0.03 (low-ratio) and 0.12 (high-ratio). Spearman correlations between estimated cell-type abundance and immune activation markers identified one significant association: in B-cell follicles, B-memory cell abundance was positively correlated with HIV DNA (ρ = 0.89, p = 0.007, q = 0.16; Fig. 5E).

## Discussion

Spatial transcriptomics of ileal and inguinal lymph node tissue from PWH on long-term ART revealed compartment-specific transcriptional programs distinguishing low- from high- CD4/CD8 ratio individuals across five anatomically and functionally distinct tissue compartments, supporting spatial heterogeneity in immune activation as a feature of persistent immune dysregulation in treated HIV infection. Low-level HIV provirus activity, epithelial barrier dysfunction with microbial translocation, and co-infections are known drivers of this persistent immune activation, yet how they interact and where they predominate within tissue compartments has remained unclear (3,8,9,13). The spatially resolved signatures identified here provide evidence of their compartment-specific footprints, mapping each contributor to distinct tissue regions rather than treating them as uniform systemic drivers. Spatial preservation of tissue architecture was critical to this resolution, enabling detection of signals that bulk or single-cell approaches would obscure, including pathway-level enrichment in the lamina propria in the absence of individually significant DEGs, or opposing ribosomal protein signatures between B-cell follicles and the T-cell zone.

Principal component analysis confirmed global expression differences between groups in four of five compartments, and the transcriptional signatures, defined by differentially expressed genes and pathway enrichment, were consistent with the known functional roles of each tissue region, a pattern additionally supported by complementary cell type composition estimates in three compartments. In the T-cell zone, these signatures were further validated by direct associations between gene expression and external measures of HIV reservoir burden and systemic immune activation.

Within the GALT, Peyer’s patches serve as the primary inductive site of mucosal immunity, where antigens are sampled and immune responses are initiated (18,19). The compartment’s transcriptional profile was consistent with this role: 57 enriched pathways spanning innate immune activation, cytokine signaling, antigen presentation, phagocytosis, oxidative stress, mitochondrial proteostasis, and translational upregulation reflect a state of persistent stimulation driven by sustained antigen exposure (19,20). Peyer’s patches were the only compartment with enrichment across all three disease branches - bacterial, parasitic, and viral infection modules, consistent with their role as the primary mucosal antigen sampling site where the broadest range of immune stimuli converge. Cell deconvolution supported this interpretation: neutrophils, NK cells, and CD4+ memory T cells were all estimated at higher proportions in low-ratio individuals, consistent with active innate and adaptive immune engagement within the compartment, while CD4+ naive T cells were estimated at higher proportions in high-ratio individuals, likely reflecting the increased peripheral blood CD4+ T cell counts in this group as the CD4/CD8 ratio in blood has shown to strongly correlated with the CD4/CD8 ratio in GALT (47). At the gene level, upregulated genes in the low-ratio group implicated altered vesicular trafficking and enhanced lymphocyte survival, while genes more highly expressed in the high-ratio group suggest greater lymphocyte proliferation and extracellular matrix activity.

Anatomically adjacent but functionally distinct, the ileal epithelium and lamina propria represent immunological compartments where luminal signals at the mucosal barrier are first sensed and then acted upon (31,35). By forming a physical barrier, the epithelium serves as the primary interface between the gut lumen and underlying tissue, acting as a regulator of intestinal immune homeostasis by actively discriminating between commensal and pathogenic bacteria through luminal antigen sampling at the epithelial surface (31,35,36,48,49). The lamina propria is also able to sample antigens from the intestinal lumen via dendritic extensions and serves as a primary effector site where activated lymphocytes carry out downstream immune responses shaped by antigens sampled from both compartments (18,19,31,36,49,50). Together they showed signatures consistent with barrier-associated stress and sustained innate immune engagement. Further studies incorporating the microbiome will also help clarify the role of microbial dysbiosis in driving inflammation from the GALT.

In the epithelium, upregulated genes in the low-ratio group collectively implicated active cellular stress and innate immune engagement at the mucosal barrier. Metallothioneins (MT1A and MT1H) involved in cellular stress response, metal ion sequestration as a defense mechanism against luminal bacteria, have previously associated with apoptosis resistance in circulating monocytes during HIV viremia (21). Also upregulated were ITLN1, a goblet cell-derived pattern recognition lectin that binds microbial glycans as part of mucosal innate defense, and BLNK, a B-cell receptor signaling adaptor whose expression in an epithelial compartment may reflect immune cell infiltration during barrier disruption (51,52). The three taken together may suggest an active stress and damage response at the mucosal surface, providing a potential mechanism for decreased epithelial barrier integrity in HIV infection. Consistent with active mucosal engagement, estimated endothelial cell fractions were higher in the low-ratio group. Intestinal endothelial cells are abundant in gut mucosal tissue, interact regularly with infiltrating immune cells, and have been shown to promote HIV infection and latent reservoir formation in CD4+ T cells in the gut microenvironment (53). Endothelial activation and vascular remodeling are recognized features of intestinal mucosal inflammation, and expansion of the endothelium through angiogenesis, mediated by inflammatory cytokines and growth factors, is a previously reported mark of active inflammatory guts diseases related to severity (54,55). Their potential contribution to barrier dysfunction and immune cell trafficking in this context warrants further investigation. Higher expression of FCMR/Toso and CCL3L3 in the high-ratio group of this compartment mirrored findings in the T-cell zone, suggesting that reduced Fas-mediated apoptosis resistance and diminished CCR5-competitive chemokine activity are shared features of epithelial barrier damage and lymphoid dysfunction (22,56,57), again providing plausible mechanisms for epithelial barrier breakdown and chronic immune activation and inflammation in PWH.

In the lamina propria, the absence of individually significant DEGs concurrent with enrichment of pathways spanning cytokine signaling, innate and adaptive immune activation, parasite phagocytosis, and scavenger receptor-mediated ligand uptake reflects a diffuse, distributed transcriptional response consistent with its role as a broadly activated effector site, notably including immunoregulatory interactions between lymphoid and non-lymphoid cells and MHC class I antigen presentation (35,49,50). ECM proteoglycan enrichment further supports ongoing structural remodeling at this site, consistent with sustained immune cell trafficking during chronic mucosal inflammation (58,59). Finally, the co-enrichment of pro-inflammatory programs alongside FCGR3A-mediated IL-10 synthesis and type 2 cytokine signaling suggests that immune activation and regulatory suppression coexist in this compartment, potentially sustaining chronic antigen exposure while impairing effective microbicidal clearance (60,61). Enrichment of IFN-α/β signaling in both the lamina propria and Peyer’s patches suggests that persistent HIV proviral antigen may be contributing to ongoing innate immune activation in the gut mucosa despite systemic viral suppression; while type I IFN responses initially limit HIV spread, chronic exposure in ART-treated individuals is associated with immune desensitization and sustained immune activation rather than effective clearance (62), thus this may lead to chronic inflammation observed in PWH.

The inguinal lymph nodes serve as both inductive and effector sites, where adaptive immune responses are initiated and effector populations are generated and maintained (30). The transcriptional signatures observed in each compartment were consistent with these specialized roles. B-cell follicle pathway enrichment was dominated by translational machinery and RNA processing in the low ratio group. It also included nutrient-sensing pathways, and a sole enriched immune pathway being regulation of signaling by CBL, a negative regulator of cytokine receptor signaling. This biosynthetically active profile, with canonical immune signaling limited to a single negative regulatory pathway, is consistent with an environment of active lymphocyte activation and differentiation driven by chronic antigen exposure in low-ratio individuals. In contrast, cell deconvolution revealed that B-memory cells were estimated at higher proportions in high-ratio individuals and showed a strong positive association with HIV DNA (ρ = 0.89, q = 0.16). This association is notable given the lower systemic immune activation profile of high-ratio individuals, and may reflect compartmentalized follicular reservoir dynamics not captured by systemic markers (24,63). B-cell follicles are established sanctuary sites for HIV where follicular CD4+ T-cell subsets are more permissive to infection and CTL surveillance is limited (63), and higher estimated Treg proportions in low-ratio individuals may reflect a more immunosuppressive follicular microenvironment, consistent with the known role of follicular regulatory T cells in modulating germinal center responses during HIV infection (64). The nature of these relationships warrants further investigation.

The T-cell zone showed the greatest number of DEGs of any compartment alongside the fewest enriched pathways, pointing toward innate immune complex clearance, lipid handling, and antigen uptake as the dominant transcriptional programs. This combination of broad gene-level divergence and narrow pathway enrichment suggests a microenvironment where diverse cellular functions co-occur under sustained inflammatory pressure without converging into a single dominant response. Notably, the T-cell zone was the only compartment in which gene expression showed significant associations with external measures of HIV reservoir burden and systemic immune activation, linking tissue-level transcriptional programs directly to systemic disease markers. Further studies delineating the HIV viral reservoir relative to T cell functions in the T-cell zone are warranted.

Despite their compartment-specific character, the signatures described above were unified by recurring molecular themes suggesting a common inflammatory infrastructure expressed differently across tissue regions, convergent mitochondrial stress and inflammasome activation, impaired immune execution despite active upstream signaling, and divergent translational reprogramming reflecting distinct metabolic demands across lymphoid microenvironments. Taken together, these data provide evidence for the mechanisms which underlie decreased epithelial barrier integrity, persistent inflammation and immune activation in HIV infection.

ORMDL3 and ARL17B were both upregulated across Peyer’s patches, B-cell follicles, and the T-cell zone. ORMDL3 overexpression induces mitochondrial fragmentation and enhances ER-mitochondria contacts, enabling NLRP3 inflammasome activation and downstream release of IL-1β and IL-18, an axis previously implicated in ulcerative colitis (65). Literature on ARL17B is scarce, though it has been identified as a candidate modulator of mitochondrial morphology whose depletion promotes mitochondrial fragmentation and reduces intracellular ATP levels (66). Additionally, ARL17 depletion impairs influenza A viral replication through reduced ATP availability and delayed nuclear export of viral ribonucleoprotein complexes, suggesting a broader role in supporting intracellular RNA-protein trafficking (67). While HIV does not utilize ribonucleoprotein complexes in the same manner as influenza A, ARL17B may nonetheless participate in trafficking pathways relevant to HIV pathogenesis given the known dependence of HIV replication on host vesicular machinery and mitochondrial bioenergetics. The association of ARL17B with HIV, particularly with higher expression in PWH with low CD4/CD8 ratios, to our knowledge has not been reported and could warrant further investigation given its association with other viral infections.

A pattern of high immune activation alongside impaired effector function was also apparent. Kinesin motor activity was the only negatively enriched pathway and found in Peyer’s patches, raising the possibility of reduced intracellular transport capacity alongside otherwise broadly activated immune programs (68). FCMR, a regulator of Fas-mediated apoptosis, was downregulated in the epithelium and T-cell zone of high-ratio individuals, consistent with prior evidence that Fas-induced apoptosis of CD4+ T cells is inversely correlated with CD4+ T cell recovery after ART (69). Further, CCL3L3, a CCR5-binding chemokine that competitively inhibits HIV entry, was downregulated across three compartments in low-ratio individuals (56,57), suggesting a relationship between increased HIV susceptibility and the compartment specific transcriptomic signatures observed between our groups.

Translational reprogramming was a third recurring theme, with divergent directionality between compartments. Ribosomal protein genes were extensively upregulated in the low-ratio group in B-cell follicles, consistent with biosynthetically active germinal center responses (70), while a distinct set of RPL and RPS genes was upregulated in the opposite direction in the high-ratio T-cell zone. Several of these ribosomal proteins, including RPL13, RPL14, RPL17, and RPL35A, have documented alterations in nucleolar abundance following HIV-1 Tat expression in T cells (71), raising the possibility of Tat-mediated contributions to divergent translational programs across lymphoid compartments. Shared enrichment of EIF2AK4-mediated amino acid deficiency sensing and selenocysteine synthesis across Peyer’s patches and B-cell follicles further indicates metabolic stress adaptation in organized lymphoid tissue (72–74).

Correlations between T-cell zone DEGs and external biomarkers, identified exclusively in this compartment, provided additional hypothesis-generating evidence. SERINC5, a host restriction factor that impairs HIV membrane fusion and entry (42–44) was inversely associated with HIV RNA and enriched in the high-ratio group, providing a plausible mechanism for differences observed in viral reservoirs.

The marker of microbial translocation and innate immune activation sCD14 showed bidirectional associations with T-cell zone DEGs across both ratio groups, with positive associations restricted to low-ratio genes including TIGIT, an inhibitory checkpoint receptor associated with T-cell exhaustion (75,76), STMN1, a regulator of microtubule dynamics linked to cellular activation (45). High-ratio genes showed inverse associations, notably FCGBP, a mucosal glycoprotein associated with pathogen exclusion and barrier defense (46), together suggesting that microbial translocation, reduced host restriction, and T-cell exhaustion represent converging features of T-cell zone dysregulation in this compartment.

This study should be interpreted in the context of several limitations. The overall cohort was modest, comprising 10 individuals, of whom eight contributed tissue for each analysis (four per ratio group), with multiple ROIs collected per individual. Tissue samples were selected based on the presence of key histological structures, which was necessary for compartment-level analysis but introduces a selection bias toward samples with more intact or active tissue architecture. Several potential confounders were not available for adjustment, including diet, lifestyle factors, and MSM status; while age and length of HIV infection were known, 55 vs. 31 years and 12 vs. 3 years respectively, formal adjustment was not performed given our limited sample size. Given the well-established influence of these factors on immune profiles, their absence constrains causal interpretation of the observed associations. Cell type deconvolution and gene-correlation analyses were evaluated at a relaxed FDR threshold of q ≤ 0.25, reflecting the exploratory nature of these analyses and the limited statistical power available at q ≤ 0.05 in this dataset; findings from these analyses should be considered hypothesis-generating rather than confirmatory. Finally, the CD4/CD8 ratio groups used to define low- and high-ratio individuals represent surrogate markers of immune dysfunction rather than directly observed clinical outcomes, and future studies with longitudinal data would allow these transcriptional signatures to be linked to specific health endpoints. Notwithstanding these constraints, the findings presented here are biologically consistent, spatially resolved, and provide a foundation for hypothesis-driven validation in larger and more diverse cohorts. Further, we defined several potential mechanisms by which chronic immune activation and inflammation may persist in HIV infection and lead to morbidities and mortality.

## Methods

### Participant enrollment and sampling

This analysis included ten PWH enrolled in a study led by the University of Minnesota (UMN) between July 2021 and December 2023. Participants were required to be ≥ 18 years of age, laboratory-confirmed for HIV-1 infection, virally suppressed on an ART regimen for >12 months with plasma HIV RNA <48 copies/mL, and within normal range for blood screen tests (complete blood count and metabolic panel). Exclusion criterion included a BMI ≥ 30 kg/m^2^, use of anticoagulants, and a history of ≥3 lymph node biopsies in the past. No participants were on antibiotics at the time of enrollment. The UMN IRB approved the study (STUDY00009216), and all participants gave informed consent using IRB-approved forms.

Colonoscopies were conducted at the Endoscopy Center of the UMN Medical Center (UMMC) under mild-to-moderate sedation and after overnight bowel preparation. Excisional inguinal LN biopsies were completed by UMN surgeons at the Clinical Research Unit (CRU) under ultrasound guidance and with administration of local anesthesia. Blood was collected in EDTA tubes prior to either colonoscopy or lymph node biopsy with instructions to fast. Within 30 minutes of collection, plasma was isolated by centrifugation and stored at -80°C.

## GeoMx slide preparation

Samples used here were selected based on the presence of key histological structures in tissue sections, specifically, Peyer’s patches, lamina propria, and intact epithelium in ileal samples, and B cell follicles and T cell zones in lymph node samples. Selected samples were then stratified into two groups by CD4/CD8 ratio, defining low-ratio and high-ratio groups as above or below the median.

Formalin-fixed, paraffin-embedded (FFPE) tissue samples were sectioned at 5-μm thickness following NanoString sample preparation guidelines. Sections first underwent antigen retrieval with a high pH EDTA based antigen retrieval buffer, followed by digestion with proteinase K to expose RNA targets for optimal probe hybridization and then fixed in 10% neutral buffered formalin. Prepared tissues were then flooded and hybridized with whole transcriptome RNA probes (v1.0, Cat # 121401102, Nanostring Technologies) attached to oligonucleotide barcodes via photocleavable linkers. Following tissue preparation, slides were stained with fluorescent visualization markers including CD45 (Cell Signaling Technology, Cat# 10143BC) and PanCK (NovusBio, Cat# NBP2-33200) for ileum tissue, CD3 (RM-9101, clone SP7) and CD20 (M0755, clone L26) for lymph nodes and SYTO 13 fluorescent nucleic acid stain (ThermoFisher, Cat#S7575). Region of interest (ROI) selection was guided by fluorescent markers, spatial location, and morphological features. Peyer’s patches, B cell follicles, and T cell zone structures were selected using geometric ROIs, whereas segmented ROIs were used for epithelium (PanCK+, CD45-) and lamina propria (PanCK-, CD45+). From these ROIs, the GeoMx DSP instrument utilizes an ultraviolet laser to release oligonucleotide barcodes from the area of illumination (AOI), which are collected in 96-well plates for library preparation and sequencing.

## GeoMx library preparation and sequencing

Library preparation was performed according to the manufacturer’s instructions. Briefly, samples were evaporated overnight and resuspended in diethylpyrocarbonate (DEPC)- treated water, followed by a brief incubation at room temperature. PCR amplification was then performed to ligate Illumina adapter sequences and unique dual-index sample barcodes. Following PCR amplification, libraries were pooled and purified by two rounds of AMPure XP bead cleanup (Beckman Coulter, Cat# A63880). Pooled libraries were quantified using the Qubit dsDNA HS kit assay (Invitrogen, cat. Q32854) on a Qubit 3.0 Fluorometer (Invitrogen) followed by quality and amplicon size assessment on an Agilent TapeStation.

Sequencing read depth was calculated based on the manufacturer’s recommendation of 100 reads per square micron of Area of Illumination (AOI). Ileum and lymph node samples were library prepped and sequenced independently, targeting 3.45x10^6^ and 9.94x10^5^ total reads, respectively. Sequencing was performed on an Illumina NovaSeq S4 300 cycle flow cell generating 2 x 27 base-pair reads. In the downstream analysis below, each AOI is referred to as a segment.

## GeoMx data preprocessing

Preprocessing was performed in R (version 4.4.3) using the GeomxTools package (26), following recommended quality control workflows applied sequentially at the segment, probe, and gene level. Firstly, this included a minimum of 1,000 raw sequencing reads per segment, ≥80% trimmed and stitched reads, ≥75% aligned reads, and ≥50% sequencing saturation. Segments were further required to have ≥50 nuclei and a minimum area of 1,000 μm². Segments failing to meet these criteria were flagged and removed from downstream analysis. Negative control counts, representing the geometric mean of negative probes in the GeoMx panel that do not target mRNA, were required to be ≥3 to establish background count levels and no-template control (NTC) wells were monitored to detect contamination.

At the gene level, segments in which fewer than <5% of panel genes were detected above the limit of quantification (LOQ), as defined by the negative probes, were removed. Genes detected above the LOQ in ≥10% of segments were retained for analysis, and genes identified as local outliers within their respective segments were excluded. After filtering, the number of genes retained from the complete set of 18,677 was 7,604 in the epithelium, 4,415 in the lamina propria, 11,578 in Peyer’s patches, 10,398 in the B cell follicles and 11,222 T cell zone of lymph structures.

Normalization and batch correction were further performed using the standR package (77). Briefly, gene expression data were normalized using the Trimmed Mean of M-values (TMM) method to account for differences in library size and RNA composition across samples (78). Batch effects across slides were corrected using the Remove Unwanted Variation (RUV4) method implemented in the geomxBatchCorrection function (79).

## Serological assays

Plasma IL-6 concentrations were measured by Luminex multiplex immunoassay (MILLIPLEX, Cat# HSTCMAG-28SK). Markers of gut barrier integrity and microbial translocation were quantified by ELISA, including intestinal fatty acid-binding protein (I-FABP; R&D Systems, Cat# DFBP20), lipopolysaccharide-binding protein (LBP; Abcam, Cat# Ab279407), and soluble CD14 (sCD14; R&D Systems, Cat# QK383).

## HIV reservoir measurements

The HIV reservoir was quantified in 4% paraformaldehyde-fixed lymph node biopsies as previously described (80–83). Briefly, five to ten 5 µm sections separated by 20 µm intervals were analyzed by RNAscope 2.5 (Advanced Cell Diagnostics) using in situ hybridization with HIV-specific probes targeting Clade B viruses (RNA antisense probe, Cat# 416111; DNA sense probe, Cat# 425531). Quantitative image analysis was used to determine the number of HIV RNA+ and DNA+ cells per unit tissue area.

## Statistical analysis

Principal component analysis (PCA) was used to assess global gene expression differences between groups using RUV4 normalized expression data. Differential expression analysis was performed using the limma-voom pipeline (84,85) on normalized and batch-corrected gene abundances with repeated measures accounted for using a random effect for patient ID and genes considered significant at FDR q ≤ 0.05. Gene set enrichment analysis (GSEA) was performed using preranked gene lists based on a composite score (log_2_FC × −log10(p-value)) (86), with enrichment tested against Reactome pathways (87), with significance defined as FDR q ≤ 0.05 and pathway relationships interpreted using the Reactome hierarchical structure. Cell type composition was estimated using SpatialDecon with the safeTME reference matrix (88), and between-group differences in cell type proportions were tested using the propeller framework from the speckle package (89), with significance defined as nominal p-value ≤ 0.05 and FDR q ≤ 0.25. Spearman correlations were performed between TMM-normalized gene expression levels (averaged across repeated measures) and selected external measurements including plasma IL-6, gut damage biomarkers (LBP, I-FABP, sCD14), and IHC quantified HIV reservoir measures in lymph nodes (viral RNA and DNA). Significant correlations were defined by nominal p ≤ 0.05 and FDR q ≤ 0.25. This analysis was also repeated using differentially abundant cell type proportions within compartments. Gene functions and protein annotations referenced throughout were obtained from NCBI Gene and GeneCards, and HIV-host interaction data were obtained from the HIV-1 Human Interaction Database (39–41).

## Study approval

All participants gave written informed consent using IRB-approved forms. The UMN Institutional Review Board approved this study (STUDY00009216).

## Data availability

Sequencing data available at NCBI under Sequence Read Archive submission SUB16438386. All code used in this analysis available from the corresponding authors upon reasonable request.

## Contributions

Conceptualization: NRK, TWS, CMB

Supervision: NRK, TWS, JA, KE, CMB

Funding acquisition: NRK, TWS, CMB

Investigation: MB, CMB, GW, JA, TS, ES

Formal analysis: MB

Visualization: MB

Writing – original draft: MB

Writing – review & editing: MB, CMB, NRK, MG

## Acknowledgements

We are deeply grateful to the participants of this study. We would also like to thank Dr. Alexander Khoruts who performed the colonoscopies, and Dr. Greg Beilman and Dr. Jeffrey Chipman who performed the lymph node biopsies. Thank you to the UMN University Imaging Centers, Erin Hudson, and Dr. Fernanda Rodriguez for assistance with the GeoMx data generation.

## Abbreviations

AOI: Area of Illumination
ART: Antiretroviral therapy
BCR: B cell receptor
BMI: Body mass index
CD: Cluster of differentiation
CMV: Cytomegalovirus
CRP: C-reactive protein
CRU: Clinical Research Unit
DAVID: Database for Annotation, Visualization, and Integrated Discovery
DEGs: Differentially expressed genes
DEPC: Diethylpyrocarbonate
DNA: Deoxyribonucleic acid
DSP: Digital Spatial Profiler
EBV: Epstein-Barr virus
ECM: Extracellular matrix
EDTA: Ethylenediaminetetraacetic acid
ELISA: Enzyme-linked immunosorbent assay
ER: Endoplasmic reticulum
FDR: False discovery rate
FFPE: Formalin-fixed, paraffin-embedded
GALT: Gut-associated lymphoid tissue
GI: Gastrointestinal
GSEA: Gene set enrichment analysis
HIV: Human immunodeficiency virus
HPCs: Hematopoietic progenitor cells
IFN: Interferon
I-FABP: Intestinal fatty acid-binding protein
IHC: Immunohistochemistry
IL: Interleukin
IRB: Institutional Review Board
LBP: Lipopolysaccharide-binding protein
LOQ: Limit of quantification
M.tb: Mycobacterium tuberculosis
MHC: Major histocompatibility complex
MSM: Men who have sex with men
NES: Normalized enrichment score
NK: Natural killer (cells)
NMD: Nonsense-mediated decay
NTC: No-template control
PCA: Principal component analysis
PCR: Polymerase chain reaction
PD: Percent difference
pDCs: Plasmacytoid dendritic cells
PERMANOVA: Permutational multivariate analysis of variance
PWH: People with HIV
RNA: Ribonucleic acid
ROI: Region of interest
SARS-CoV: Severe acute respiratory syndrome coronavirus
sCD14: Soluble CD14
TMM: Trimmed Mean of M-values
Tregs: Regulatory T cells
UMMC: University of Minnesota Medical Center
UMN: University of Minnesota

